# Distribution of *bla*_CTX-M_-gene variants in *E. coli* from different origins in Ecuador

**DOI:** 10.1101/2023.03.15.532797

**Authors:** Xavier Valenzuela, Hayden Hedman, Alma Villagomez, Paul Cardenas, Joseph N. S. Eisenberg, Karen Levy, Lixin Zhang, Gabriel Trueba

## Abstract

The increasing abundance of extended spectrum β-lactamase (ESBL) genes in *E. coli*, and other commensal and pathogenic bacteria, endangers the utility of third or more recent generation cephalosporins, which are major tools for fighting deadly infections. The role of domestic animals in the transmission of ESBL carrying bacteria has been recognized, especially in low- and middle-income countries, however the horizontal gene transfer of these genes is difficult to assess. Here we investigate *bla*_CTX-M_ gene diversity (and flanking nucleotide sequences) in *E. coli* from chicken and humans, in an Ecuadorian rural community and from chickens in another location in Ecuador. The *bla*_CTX-M_ associated sequences in isolates from humans and chickens in the same remote community showed greater similarity from those found in *E. coli* in a chicken industrial operation 200 km away. Our study may provide evidence of *bla*_CTX-M_ transfer between chickens and humans in the community.

## 1. Introduction

Antimicrobial resistance is recognized as one of the top 10 global public health threats facing humanity [1, 2, 3], with an estimated 700,000 annual deaths due to antibiotic-resistant infections, and predictions of 10 million deaths by 2050 [1, 3]. Every year, reports on multi-resistant microorganisms increase in different countries, indicating that we are heading into a post-antimicrobial era in which common infections and minor wounds can cause death [4]. Different prevention and control measures have been proposed to mitigate this problem but with minimal success [1, 2, 5].

Most antimicrobials are administered to animals raised for food, and antimicrobial use in animals is projected to increase to over 100,000 tons by 2030, as global demand for animal protein increases [6]. In Ecuador and in other low- and middle-income countries (LMICs), increased small-scale farming has resulted in greater use of antibiotics as growth promoters (and prophylactics), selecting antimicrobial-resistant bacteria which colonizes different hosts and contribute to the dissemination of antimicrobial genes [7, 8 19, 20].

One of the main antimicrobial resistance genes codes for extended spectrum β-lactamases (ESBLs) which are a rapidly evolving group of enzymes able to hydrolyze third generation cephalosporins, one of the most widely used antibiotics in hospitals and intensive care units [10]. These genes are carried on mobile genetic elements (MGE) which facilitate their spread between bacterial species belonging to different animal microbiomes [9].

*Escherichia coli* is considered one of the most serious threats in the antimicrobial resistance crisis [11] and it belongs to a type of intestinal bacteria that transfers the most amongst different animal hosts [12]. The *bla*_CTX-M_ is the most widely distributed ESBL gene, even though it started to disseminate in *E. coli* (from the environmental bacteria *Kluyvera* spp.) as recently as the 1990s [13,14,15]. The *bla*_CTX-M_ genes have displaced other ESBLs genes in Enterobacteriaceae, maybe due to highly active MGEs (transposons and plasmids) and successful bacterial clones [13, 16, 17]. Currently, there are more than 234 *bla*_CTX-M_ allelic variants in *E. coli* [18] and only a small number are successful.

In *E. coli*, the dissemination of *bla*_CTX-M_ genes occurs clonally and through horizontal gene transfer (HGT). Evidence of antimicrobial-resistant bacteria transmission, from domestic animals to humans, is difficult to obtain; clonal transmission between domestic animals and humans requires the search for genetically identical isolates from a large diversity of *E. coli* lineages [19, 20]. Evidence of HGT requires finding identical plasmids, with the complication that plasmids are diverse, plasmid DNA could be constantly rearranged [21], and resistance genes are usually in highly active transposons (moving back and forth among plasmids, from plasmid to chromosome, or to other MGEs) [22, 23, 24, 27].

Some reports showing different *bla*_CTX-M_ allelic variants in *E. coli* isolates from humans and domestic animals, concluded that role of domestic animals in the current antimicrobial crisis may have been overestimated [25, 26]. However, other studies using isolates with closer spatial-temporal relationships, showed similar allelic variants in isolates from humans and domestic animals [19, 20]. Here we analyzed the distribution of *bla*_CTX-M_ allelic variants and flanking sequences (in conjugable plasmids) from human and chicken isolates in the same community from 2010-2017 and compared with similar *bla*_CTX-M_ allelic variants in *E. coli* isolated from a chicken industrial operation 200 km away, during overlapping periods.

## 2. Materials and methods

### 2.1. Isolates

We analyzed 107 *E. coli* 3^rth^ generation cephalosporin resistant isolates (from a total of 4,518 *E. coli* isolates analyzed): 38 (35.5%) from humans (in a rural community) and 69 (64.5%) from chickens of which 26 (37.7%) from the same rural community and 43 (62.3%) were from an industrial operation (Table 2). The isolates were obtained from previous studies (from humans and chickens) in rural communities in northern Coastal Ecuador during 2017 [7, 8], in 2009 [28], and 2010-2013 [29] and from chickens from an industrial operation (200 km apart from the remote community) during 2010. In these studies, *E. coli* was isolated from fecal samples by streaking on MacConkey Agar and incubating for 24 hours at 37°C. Lactose fermenting colonies were tested for β-glucuronidase production, as described previously [7, 8]. Resistance profiles were assessed using the Kirby-Bauer disk diffusion method following Clinical and Laboratory Standards Institute (CLSI) recommendations. We used *E. coli* ATCC 25922, *Staphylococcus aureus* ATCC 25923, and *Pseudomonas aeruginosa* ATCC 27853 as controls for antimicrobial susceptibility test. All *E. coli* were stored frozen at −80°C until analyzed. Out of 4,518 *E. coli* isolates, we identified 107 with phenotypic third-generation cephalosporin-resistance (TGCR). All the protocols were approved by Universidad San Francisco de Quito’s Bioethics committee and University of Michigan’s IRB.

### 2.2. Conjugation experiments

To corroborate that the genes were in conjugable plasmids, we conducted conjugation experiments to identify the *bla*_CTX-M_-genes that had the potential to be horizontally transferred by plasmids. Conjugation experiments were performed in Lysogeny Broth (LB) with *E. coli* J53Az^r^ (resistant to sodium azide and susceptible to cefotaxime) as the recipient, and all of our *E. coli* TGCR isolates as donors [30]. We cultured donor and recipient isolates in tubes with 5 ml of LB incubated for 12 hours at 37°C, to get cells in a logarithmic growth phase. We then added 0.5 ml of each tube to 4 ml of fresh LB and incubated for 16 hours at 37°C without shaking. Transconjugants were selected using selective culture media made of Trypticase soy agar (TSA) supplied with sodium azide (200 μg/ml) and cefotaxime (1mg/ml). All transconjugants were stored frozen at−80 °C.

### 2.3. Polymerase chain reaction [PCR] and DNA sequencing

Genetic materials from all 107 *E. coli* TGCR isolates were extracted using DNAzol® (Invitrogen™, USA) following manufacturer protocol and recommendations. We performed conventional PCR to identify samples carrying *bla*_CTX-M_-gene (PCR1); to identify the sequences upstream from the *bla*_CTX-M_-gene (PCR2), and to identify sequences downstream from the *bla*_CTX-M_-(PCR3).

PCR1 was carried out using degenerate primers [31]. PCR2 used the *bla*_CTX-M_-gene and IS*Ecp1* sequences [31]. Both sequences are often in close proximity. PCR3 was designed to obtain complete coding sequences of *bla*_CTX-M_-genes and the downstream sequences. We used degenerate primers of *bla*_CTX-M_- and *orf477* [32]. All amplicons obtained were sequenced. Primers used in this study are listed in Table 1 and the locations of the flanking regions are shown in Figure 1. Accession numbers of the sequences are in supplemental material, Table S1. Amplicons were sequenced using Sanger’s method at Research Technology Support Facility, Michigan State University.

**Table 1.**
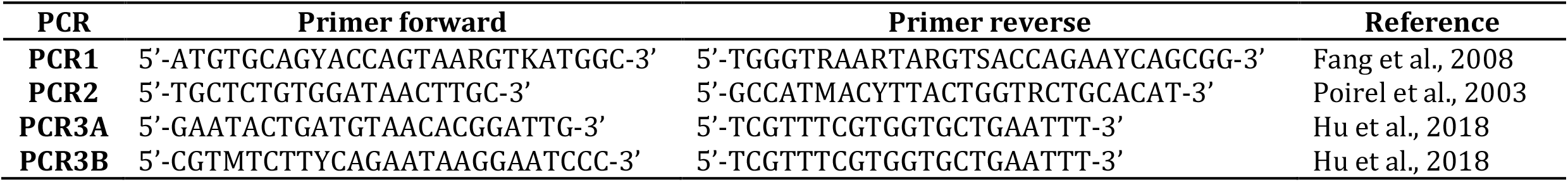
Sequences of primers used in PCR experiments in this study. PCR experiment described in the text.

**Figure 1.**
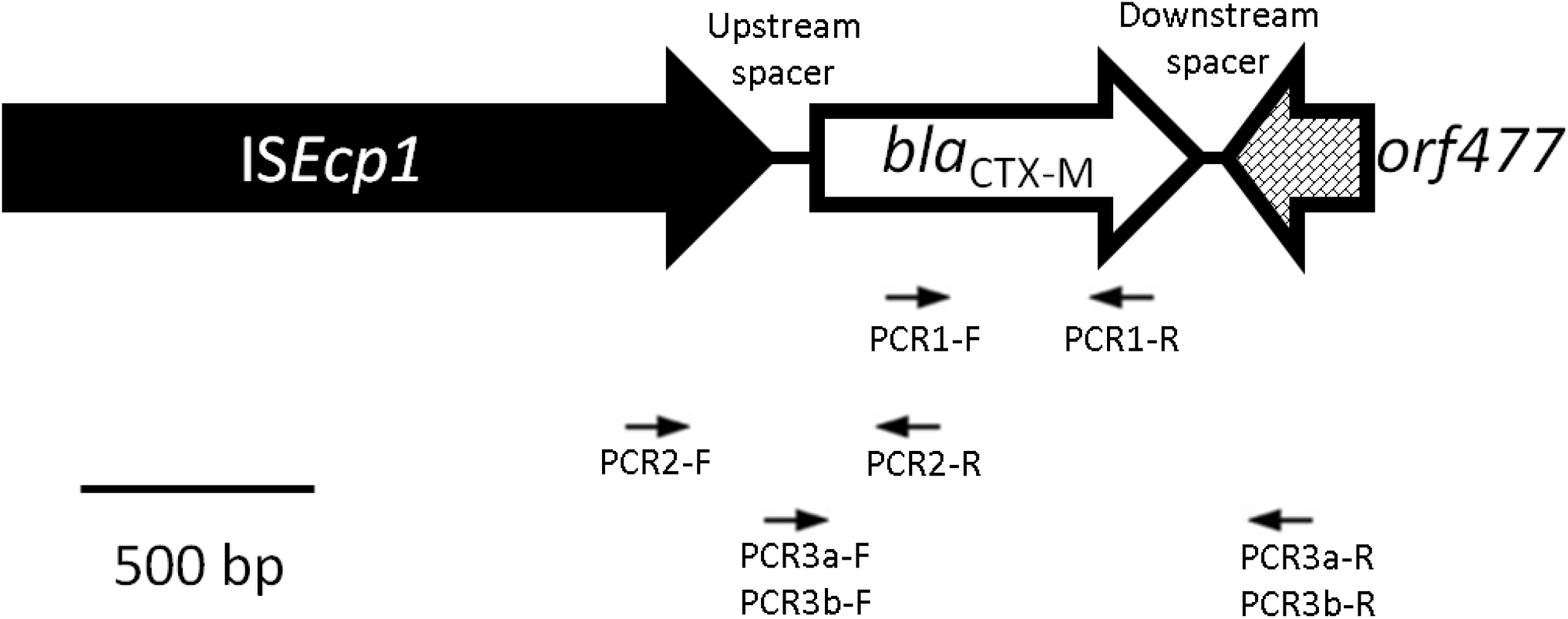
Schematic representation of usual orientation of the *bla*_CTX-M_-genes. Here we see the transposon IS*Ecp1*[Insertion Sequence] upstream from the *bla*_CTX-M_-gene, and the *orf477* gene in the downstream region. Arrows indicates the primers used in this study.

### 2.4. bla_CTX-M_-variant distribution analysis

We assumed that an evidence of gene transfer in the local community were the presence of isolates with: 1) same *bla*_CTX-M_-allele variant; 2) same number of nucleotides and same nucleotide sequences between the *bla*_CTX-M_-gene and the transposable element; 3) had the same transposon (or Insertion Sequence); and 4) had the same downstream gene (Figure 1).

### 2.5. Statistical analysis

We used R statistical software to run principal component analysis (PCA) and ANOVA to evaluate the relationship between *bla*_CTX-M_ allele variants (from humans’ and chickens’ *E. coli*) in the community and those from chickens in the industrial operation. We used GraphPrim 9.5 for the graphic representation.

## 3. Results

### 3.1. *Nucleotide sequence analysis of bla*_CTX-M_ *genes and flanking regions*

We obtained 102 transconjugants (from isolates carrying *bla*_CTX-M_ genes); 37 (36.3%) from humans and 65 (63.7%) from chickens of which 40 (61.5%) were from industrial operation and 25 (38.5%) were from the rural community (Table 2).

**Table 2.** Characteristics of *E. coli* isolates from 94 human and chicken fecal isolates, including year sample was collected, the variant number for the *bla*_CTX-M_-gene, spacer [number of base pairs between the *bla*_CTX-M_-gene and the downstream transposon], Insertion Sequence [IS], and the gene that is downstream from the *bla*_CTX-M_-gene. The sequences are ordered according to the type of *bla*_CTX-M_-variant.

| Source | Number isolates | Year | *bla*<sub>CTX-M</sub>- variant | Upstream Spacer | IS [transposon] | Down-stream Sequences |
| --- | --- | --- | --- | --- | --- | --- |
| Chicken Community | 1 | 2017 | 15 | 127 bp | ISEcp1 | N/A |
| Chicken Community | 1 | 2017 | 15 | 127 bp | ISEcp1 | ORF477 |
| Chicken Community | 11 | 2017 | 55 | 127 bp | ISEcp1 | ORF477 |
| Chicken Community | 4 | 2017 | 64 | 45 bp | ISEcp1 | N/A |
| Chicken Community | 3 | 2017 | 64 | 45 bp | ISEcp1 | ORF477 |
| Chicken Community | 4 | 2017 | 65 | 42 bp | ISEcp1 | N/A |
| Chicken industrial | 28 | 2010 | 2 | N/A | N/A | N/A |
| Chicken industrial | 5 | 2010 | 8 | N/A | N/A | N/A |
| Chicken industrial | 1 | 2010 | 12 | 48 bp | ISEcp1 | N/A |
| Chicken industrial | 1 | 2010 | 12 | 48 bp | ISEcp1 | ORF477 |
| Chicken industrial | 4 | 2010 | 14 | 42 bp | ISEcp1 | N/A |
| Human Community | 2 | 2010-2013 | 15 | 48 bp | ISEcp1 | N/A |
| Human Community | 3 | 2010-2013 | 2 | N/A | N/A | N/A |
| Human Community | 4 | 2010-2013 | 9 | 42 bp | ISEcp1 | N/A |
| Human Community | 1 | 2010-2013 | 137/15 | 42 bp | ISEcp1 | N/A |
| Human Community | 2 | 2010-2013 | 14 | 42 bp | ISEcp1 | N/A |
| Human Community | 1 | 2010-2013 | 14 | N/A | ISEcp1 | N/A |
| Human Community | 1 | 2010-2013 | 15 | N/A | ISEcp1 | N/A |
| Human Community | 1 | 2010-2013 | 64 | 48 bp | ISEcp1 | N/A |
| Human Community | 1 | 2010-2013 | 65 | 42 bp | ISEcp1 | N/A |
| Human Community | 1 | 2017 | 1 | 80 bp | ISEcp1 | N/A |
| Human Community | 1 | 2017 | 1 | N/A | N/A | N/A |
| Human Community | 1 | 2017 | 9 | 42 bp | ISEcp1 | N/A |
| Human Community | 3 | 2017 | 15 | 48 bp | ISEcp1 | N/A |
| Human Community | 1 | 2017 | 15 | 48 bp | ISEcp1 | ORF477 |
| Human Community | 1 | 2017 | 15 | N/A | ISEcp1 | N/A |
| Human Community | 1 | 2017 | 15 | N/A | N/A | N/A |
| Human Community | 4 | 2017 | 27 | 42 bp | ISEcp1 | N/A |
| Human Community | 1 | 2017 | 55 | 127 bp | ISEcp1 | N/A |
| Human Community | 1 | 2017 | 64 | 45 bp | ISEcp1 | ORF477 |

We obtained PCR1 amplicons from all transconjugants, however only 94 produced clean DNA sequences. All *bla*_CTX-M_ genes found in isolates from chickens in the rural community were also present in isolates from humans of the same community (*bla*_CTX-M_-15, *bla*_CTX-M_-27, *bla*_CTX-M_-55, *bla*_CTX-M_-64, *bla*_CTX-M_-65); none of these variants were present in isolates from the chicken industrial operation. Isolates from humans in the community shared only 2 variants with chickens in the industrial operation (*bla*_CTX-M_-2 and *bla*_CTX-M_-14) (Figure 3). The prevalence of some shared variants was different for humans and chickens in the community: *bla*_CTX-M_-15 was present in 9 human and 2 chicken isolates, *bla*_CTX-M_-55 was present in 1 human and 11 chicken isolates; *bla*_CTX-M_-64 was present in 2 human and 7 chicken isolates. The prevalence of the shared *bla*_CTX-M_-2 gene was different for isolates from humans and chickens in the industrial operation (3 and 28 respectively). Three variants (*bla*_CTX-M_-1, *bla*_CTX-M_-9, *bla*CTX-M-137/15) were present only in human isolates and 2 variants (*bla*_CTX-M_-8, *bla*_CTX-M_-12) were present only in isolates from chickens in the industrial operation. Principal component analysis showed that *bla*_CTX-M_ from the community (chickens and humans) were significantly closer that those from the industrial operation (Figure S1)

In 51 *bla*_CTX-M_ positive isolates, we were able to detect the transposable element (IS*Ecp1*), and the spacer sequences (between the *bla*_CTX-M_ and IS*Ecp1*) (Table 2). The isolates that have the same *bla*_CTX-M_ seemed to share the same spacer size except for *bla*_CTX-M_-15 (2 isolates had a 127bp spacer, while 6 isolates had a 48bp spacer) and *bla*_CTX-M_-65 (1 isolate had 48bp while 8 isolates had 45 bp) (Table 2). In 18 isolates we identified the *orf477* downstream *bla*_CTX-M_ gene, however we did not see an association with any source of samples. We were able to identify a possible recombinant of *bla*_CTX-M_-15 and *bla*_CTX-M_-137 in one isolate (Figure 2). All accession numbers for these allelic variants are listed in Table S1.

**Figure 2.**
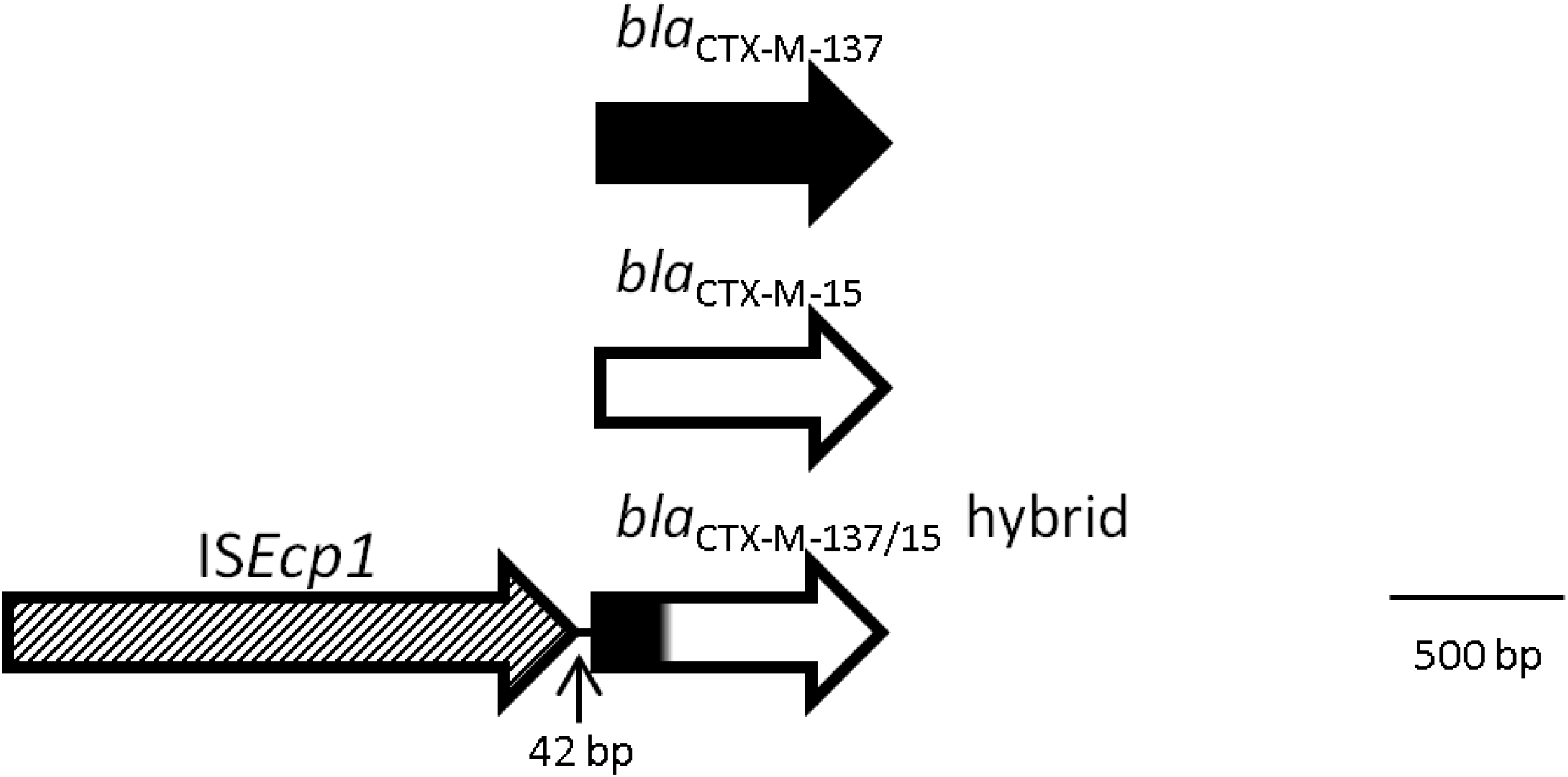
Recombination scheme between bla_CTX-M_-137 and *bla*_CTX-M_-15, sample isolated from human feces, GenBank accession number MZ314224.

## 4. Discussion

The manuscript shows the distribution of *bla*_CTX-M_ genes in *E. coli* isolates from 2 different locations in Ecuador from 2010 to 2017. All the *bla*_CTX-M_ variants (*bla*_CTX-M_-15, *bla*_CTX-M_-27, *bla*_CTX-M_-55, *bla*_CTX-M_-64, and *bla*_CTX-M_-65) and almost all flanking sequences (spacer and transposons) found in chicken isolates from the community were also found in human isolates from the same community. Contrastingly, no *bla*_CTX-M_ variants were shared between chicken’s isolates in the community and those from the industrial operation 200 km apart (even in isolates from the same year), (Figure 3). None of the *bla*_CTX-M_ variants in community’s chickens were found in the industrial chicken isolates (Table 2 and Figure 3). The PCA and ANOVA tests showed that *bla*_CTX-M_ alleles (from human and chickens) in the community are significantly closer than those from the chicken industrial operation (p=<0.0001) (Figure S1). We suggest that the distribution of *bla*_CTX-M_ variants is a supportive evidence for transference of *bla*_CTX-M_ genes among *E. coli* from chickens and humans in the same community. We did find 2 variants in isolates from chickens at the industrial operation and humans in the community (*bla*_CTX-M_-2, *bla*_CTX-M_-14), however, the proportion of isolates carrying the *bla*_CTX-M_-2 in the two species was different (15% in human’s compared to 78% in chicken’s).

**Figure 3.**
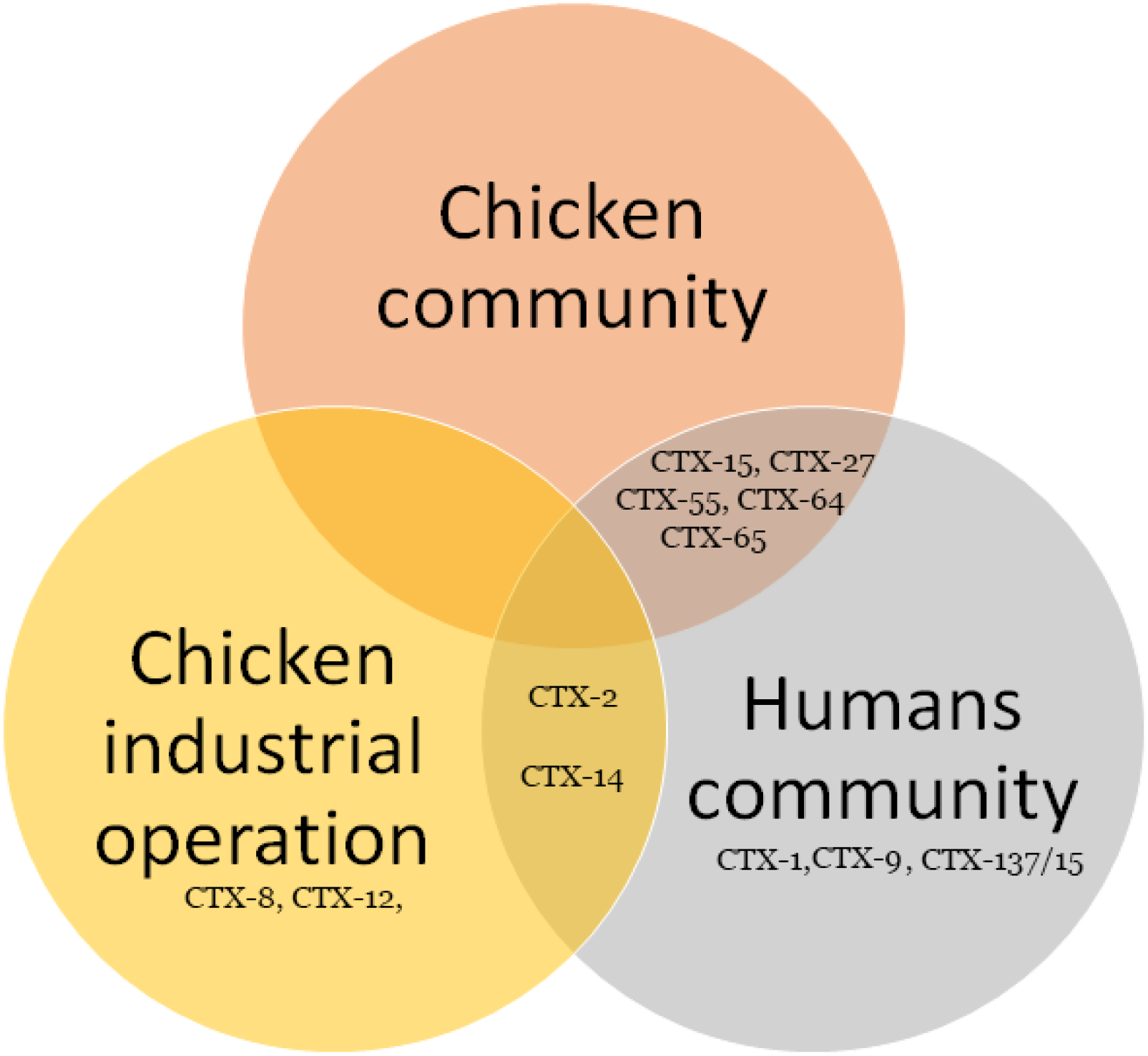
Venn Diagram showing the distribution of *bla*_CTX-M_-gene variants found in humans and chickens in different locations.

The HGT of *bla*_CTX-M_ (and other antimicrobial genes) is complex; these *bla*_CTX-M_ genes (and flanking sequences) are associated with transposable elements (such as IS*Epc1* or IS*26*) and promote high frequency transposition among different plasmids and chromosome [41, 42, 43, 44]. The transposition of genes among plasmids increases the possibilities of HGT [44], however the evidence of this transmission is difficult to detect (due to transposition), even with the use of whole genome sequencing [45]. We argue that genes associated with transposable elements and plasmids have been involved in the transference of these *bla*_CTX-M_ genes among *E. coli* in the in the community.

We did find difference in *bla*_CTX-M_ genes (and adjacent sequences) associated with different animal species; some *bla*_CTX-M_ variants (*bla*_CTX-M_-1, *bla*_CTX-M_-9) were found only in human isolates and not in chickens in the community. Additionally, spacers flanking *bla*_CTX-M_-15 in human and chicken isolates (in the same community) were different (Table 2); we also found different proportion of strains carrying same variants: human isolates had more *bla*_CTX-M_-15 than chickens in the same community (45%; n=9 vs. 8%; n=2); chicken isolates had more *bla*_CTX-M_-55 than human isolates in same community (45%; n=11 vs. 5%, n=1). These results suggest that transmission of these genes occurs more frequently among *E. coli* associated with humans or among *E. coli* associated chickens, and it may be less frequent among *E. coli* from different animal species; greater interaction with members of the same animal species may contribute to the transference of bacteria and bacterial genes among these bacteria.

The distribution of variants by time was also evident as 31% of 2017 isolates had *bla*_CTX-M_-55 whereas no isolate from 2010 to 2013 had this variant; 21% of 2017 isolates (from humans and chickens) had *bla*_CTX-M_-15 compared with 5% in isolates obtained in 2010-2013; 21% of 2017 isolates had *bla*_CTX-M_-64 whereas 1.8% of isolates from 2010-2013 had this variant (Table 2). These results are also in agreement with previous research showing that *bla*_CTX-M_ variants change overtime in a given location [22]

It was surprising to encounter more *bla*_CTX-M_ variant diversity in human isolates than in chicken isolates in the community. Additionally, we found *bla*_CTX-M_-64 and *bla*_CTX-M_-65 first in humans in the community in 2010-2013 and later in chickens in the community in 2017 (Table 2). It is tempting to speculate that these results indicate that humans may be the source of some of these gene variants found in *E. coli* from chickens. This transmission may be associated with low sanitary infrastructure in this community [33], allowing the chickens to become colonized with *E. coli* from humans. Other researchers have presented evidence of ARG transmission between *E. coli* from humans and domestic animals due to a shared environment, direct contact between humans and domestic animals, or ingestion of food contaminated with *E. coli* [7, 8, 19, 20].

Recently in Ecuador, *bla*_CTX-M_-14, *bla*_CTX-M_-15, *bla*_CTX-M_-55, and *bla*_CTX-M_-65 variants have been reported in clinical isolates [35-37]. Studies in other Ecuadorian rural communities have shown that humans and domestic animals in the same communities share *E. coli* with *bla*CTX-M-2, *bla*_CTX-M_-14, *bla*_CTX-M_-15, *bla*_CTX-M_-55, and *bla*_CTX-M_-65, suggesting a mechanism of spread (clonal and HGT of *bla*_CTX-M_ genes) between humans and domestic animals [20].

Previous research has described the close relationship between the IS*Ecp1* (and other insertion sequences) with *bla*_CTX-M_ genes [13, 38, 39]. These insertion sequences are thought to have been involved in the original mobilization *bla*_CTX-M_ genes from the bacteria *Kluyvera* spp. [13]. Most of the members of the *bla*_CTX-M_ gene family have been associated with IS*Ecp1* and five different DNA spacer sequences between the insertion sequence and the *bla*_CTX-M_ gene [13]. The length of the spacer sequence is often associated with specific a *bla*_CTX-M_ gene [40] and differences in the location of the IS*Ecp1* (Table 2) indicate frequent transposable activity, something that has been observed previously [23].

We also detected a possible recombinant of *bla*_CTX-M_-137 and *bla*_CTX-M_-15 in one isolate, being the first 218 nucleotides form *bla*_CTX-M_-137 and the rest from *bla*_CTX-M_-15 (Figure 2). It is the first report of this allelic variant; we could not find other sequences close to it during BLAST search (accession number MZ314224). We acknowledge that the small number of samples is a major limitation of our study.

We present evidence that the combination of epidemiological data (location, time frame, animal species) and DNA sequence data form *bla*_CTX-M_ genes and flanking sequences could provide evidence of *bla*_CTX-M_ genes transfer between bacteria in humans and domestic animals inhabiting the same community. We showed that nucleotide sequences of *bla*_CTX-M_ gene and flanking regions is a very useful tool to study the transmission of these genes. This information is critical to detect sources and trends in the transmission of this ESBL genes in communities.

## 5. Conclusions

The results of our research suggest that the *bla*_CTX-M_-variants are transmitted between chickens and humans in a rural community in Ecuador. We also present indirect evidence of transmission of antimicrobial resistance genes from humans to chickens in a rural community with low sanitary infrastructure. This study supports the notion that domestic animals play an important role in the circulation of ESBL genes.

## Autor contributions

Xavier Valenzuela: DNA experiments and analysis, selection of isolates, conjugation experiments, writing the manuscript. Hayden Hedman: collected samples and obtained metadata. Alma Villagomez: bacterial isolation and antibiotic susceptibility tests. Paul Cardenas: DNA analysis, data curation, and statistical analysis. Joseph Eisenberg: writing the manuscript, writing of the grant proposal. Karen Levy: writing the manuscript, writing of the grant proposal. Lixin Zhang: conceptualization, writing the manuscript, DNA sequencing, writing of the grant proposal, Gabriel Trueba: writing of the grant proposal, writing the manuscript, conceptualization, direction of the work in the laboratory

## Declaration of competing interests

The authors declare that there are no conflicts of interest

## Acknowledgments

This research was funded by the National Institute of Allergy and Infectious Diseases, National Institutes of Health, United States of America; grants: K01 AI103544 and R01 AI050038. The authors thank Deysi Parrales and Barbara Pibaque for their help with preparation of culture media and Liseth Salinas for her suggestions for the manuscript.

## Notes

### Competing Interest Statement

The authors have declared no competing interest.

